# COVID-19-related coagulopathy – Is transferrin a missing link?

**DOI:** 10.1101/2020.06.11.147025

**Authors:** Katie-May McLaughlin, Marco Bechtel, Denisa Bojkova, Mark N. Wass, Martin Michaelis, Jindrich Cinatl

## Abstract

SARS-CoV-2 is the causative agent of COVID-19. Severe COVID-19 disease has been associated with disseminated intravascular coagulation and thrombosis, but the mechanisms underlying COVID-19-related coagulopathy remain unknown. Since the risk of severe COVID-19 disease is higher in males than in females and increases with age, we combined proteomics data from SARS-CoV-2-infected cells with human gene expression data from the Genotype-Tissue Expression (GTEx) database to identify gene products involved in coagulation that change with age, differ in their levels between females and males, and are regulated in response to SARS-CoV-2 infection. This resulted in the identification of transferrin as a candidate coagulation promoter, whose levels increases with age and are higher in males than in females and that is increased upon SARS-CoV-2 infection. A systematic investigation of gene products associated with the GO term “blood coagulation” did not reveal further high confidence candidates, which are likely to contribute to COVID-19-related coagulopathy. In conclusion, the role of transferrin should be considered in the course of COVID-19 disease and further examined in ongoing clinic-pathological investigations.

## Introduction

Severe acute respiratory syndrome coronavirus 2 (SARS-CoV-2) is the causative agent of the ongoing coronavirus disease 2019 (COVID-19) outbreak [1,2]. Severe COVID-19 disease has been associated with disseminated intravascular coagulation and thrombosis [1,3,4], but the mechanisms underlying COVID-19-related coagulopathy remain unknown. However, it is known that the risk of severe and fatal COVID-19 disease is higher in males than in females and that it increases with age [5]. Similarly, the risk of coagulation-related pathologies and thrombosis increases with age and is further enhanced in males [6]. Thus, gene products that 1) are involved in coagulation, 2) change with age, 3) differ in their levels between females and males, and 4) are regulated in response to SARS-CoV-2 infection represent candidate factors that may contribute to COVID-19-related coagulopathy and disease severity.

To identify such candidate factors that may be involved in COVID-19-related coagulopathy, we here performed a combined analysis of data from a proteomics dataset derived from SARS-CoV-2-infected cells [7] and of human gene expression data from the Genotype-Tissue Expression (GTEx) database [8].

## Results

### Transferrin may be involved in COVID-19-related coagulopathy

While analysing proteomics data derived from SARS-CoV-2-infected cells [7], we found transferrin to be upregulated in SARS-CoV-2-infected cells (Figure 1A). Transferrin has been shown to increase coagulation by interfering with antithrombin/ SERPINC1-mediated inhibition of coagulation proteases including thrombin and factor XIIa [9]. Hence, there might be a link between transferrin levels and coagulation in COVID-19 patients.

**Figure 1.**
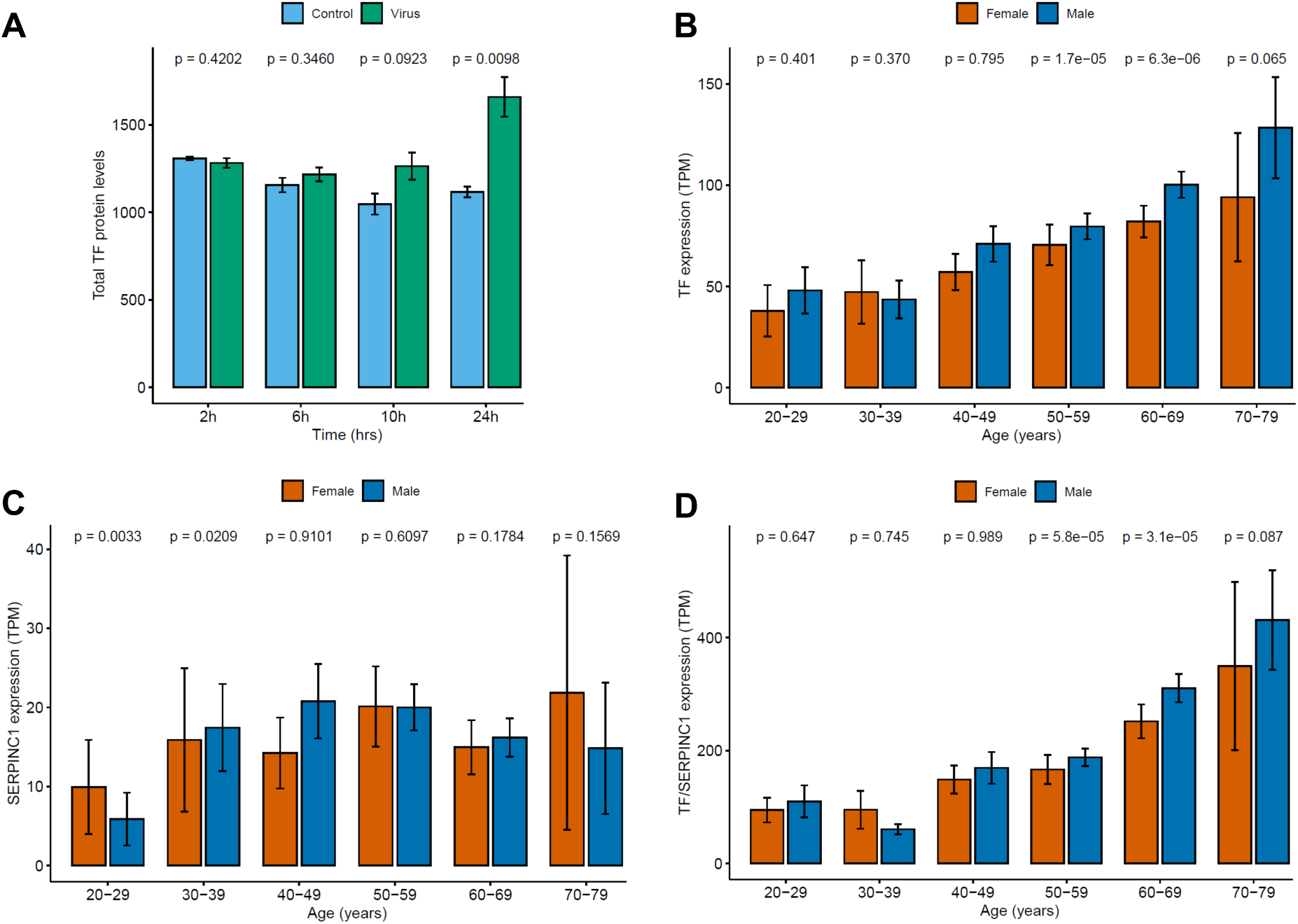
A) Mean TF protein abundance in uninfected (control) and SARS-CoV-2-infected (virus) Caco-2 cells. P-values are the result of a two-sided Student’s t-test. B) Mean TF expression (TPM) in females and males across six age groups. P-values are calculated using the Wilcoxon rank sum test for independent groups. C) Mean antithrombin (SERPINC1) expression (TPM) in females and males across six age groups. P-values are calculated using the Wilcoxon rank sum test for independent groups. D) Ratio of TF/SERPINC1 expression (TPM) in females and males across six age groups. P-values are calculated using the Wilcoxon rank sum test for independent groups.

Next, we analysed transferrin expression in the Genotype-Tissue Expression (GTEx) database [8]. We found that transferrin expression increased with age and was higher in males than in females (Figure 1B). In contrast, expression of its antagonist antithrombin did not increase with age and was similar in females and males (Figure 1C). Thus, the transferrin/ antithrombin ratio increases with age and is higher in males than in females (Figure 1D). This correlates with the risk of severe and fatal COVID-19 disease, which is higher in males than in females and also increases with age [5]. Hence, an increased transferrin/ antithrombin ratio may contribute to COVID-19-related coagulopathy and more severe disease in older patients, in particular in males.

### Analysis of genes associated with the GO term “blood coagulation” (GO:0007596) does not reveal further candidates likely to be involved in COVID-19-related coagulopathy

However, many factors are involved in coagulation and may, hence, contribute to COVID-19-related coagulopathy. Thus, we next used our approach to systematically investigate 335 gene products, which are associated with the GO term “blood coagulation” (GO:0007596) in AmiGO 2 [10]. Using GTEx, we identified 256 coagulation-associated genes, that are differently expressed between females and males (Table S1) and 237 genes whose expression changed with age (Table S2). The list of 50 overlapping genes contained many genes, whose products may be involved in coagulation under certain circumstances but are not necessarily core players directly involved in the actual coagulation process, for example members of major signalling cascades such as PI3K or MAPK signalling. Hence, the functions of these genes were manually annotated for their potential relevance as direct promoters or inhibitors of coagulation (Table S3). The expression of these 50 genes was then further analysed in the proteomics dataset comparing SARS-CoV-2-infected and non-infected cells [7], resulting in a set of nine significantly differentially expressed coagulation-associated genes (Table 1). When we combined the available data from all three datasets together with the role of the candidate factors in coagulation (promoter or inhibitor), however, we did not identify a high-confidence candidate factor (Table 1). There was neither a coagulation promoter that would be upregulated by SARS-CoV-2 infection, increase in expression with age, and display higher expression in males than in females, nor a coagulation inhibitor that would be downregulated by SARS-CoV-2 infection, decrease in expression with age, and display higher expression in females than in males (Table 1, Suppl. Figure 1).

**Table 1.** Genes with a role in coagulation that are differentially expressed between females and males (Table S1), whose expression correlates with age (Table S2), and that are differentially expressed in SARS-CoV-2-infected cells relative to control. Gene products anticipated to be of relevance would be 1) coagulation promoters that display higher expression in males than in females (FvM low), increase with age, and display high protein levels in virus-infected cells or 2) coagulation inhibitors that display higher expression in females than in males (FvM high), decrease with age, and display low protein levels in virus-infected cells.

| Gene | Coagulation | FvM | Age | Virus (p-value<sup>1</sup>) |
| --- | --- | --- | --- | --- |
| ANO6 | promoter | high | decrease | high (0.008) |
| ANXA2 | inhibitor | high | decrease | high (0.001) |
| C1QBP | promoter | high | decrease | high (0.013) |
| CEACAM1 | promoter | high | decrease | high (0.045) |
| F10 | promoter | high | decrease | low (0.044) |
| F11R | promoter | high | decrease | low (0.020) |
| GGCX | promoter | high | decrease | low (0.002) |
| MYH9 | promoter | high | decrease | high (0.004) |
| PLA2G4A | promoter | high | decrease | high (0.017) |

<sup>1</sup> determined by two-sided Student's t-test

## Discussion

Severe COVID-19 disease is associated with intravascular coagulation and thrombosis (COVID-19-related coagulopathy) [1,3,4]. The underlying mechanisms remain unclear. Our combined analysis of proteomics data derived from SARS-CoV-2-infected cells [7] and gender- and age-dependent gene expression data derived from the GTEx dataset identified transferrin as a candidate factor, which may be involved in COVID-19-related coagulopathy. Although transferrin is an iron carrier protein that circulates and delivers iron to cells via transferrin receptor binding followed by receptor-mediated endocytosis [11,12], it promotes coagulation by iron-independent mechanisms [9]. A systematic investigation of gene products associated with the GO term “blood coagulation” did not reveal further high confidence candidates, which are likely to contribute to COVID-19-related coagulopathy.

Transferrin is primarily produced in the liver, but also in other tissues [11-18]. Hence, (SARS-CoV-2-induced) locally produced transferrin may contribute to COVID-19 pathology, even independently circulating transferrin levels. For example, transferrin is produced in the brain [18], and high transferrin levels have been associated with hypercoagulability and ischemic stroke [19]. Stroke is a significant complication in COVID-19 [20], and much more common than in influenza patients [21]. Both ischemic and haemorrhagic strokes are observed in COVID-19 patients [20]. Notably, transferrin may not only contribute to ischemic strokes via inducing coagulation [19], it may also increase the brain injury associated with haemorrhagic strokes by facilitating cellular iron uptake [22]. High transferrin levels have also been associated with diabetes and metabolic syndrome [17,23-25], which are known risk factors for severe COVID-19 disease [26-28].

In conclusion, the role of transferrin in the course of COVID-19 should be considered and further examined in ongoing clinic-pathological investigations.

## Methods

### Data acquisition

Genes associated with the GO term “Blood Coagulation” (GO:0007596) were identified using the online database AmiGO 2 [10]. This generated a list of 335 unique genes annotated with 23 unique terms (including “Blood Coagulation” and 22 child terms) for further analysis.

Gene expression data (transcripts per million, TPM) and clinical data for 980 individuals (17,382 samples from 30 tissues) were downloaded from the GTEx Portal [https://www.gtexportal.org/home/datasets; GTEx Project, version 8]. We also used normalised protein abundance data from a recent publication [7] in which protein abundance in uninfected and SARS-CoV-2-infected Caco-2 cells was quantified. Data were subsequently normalised using summed intensity normalisation for sample loading, followed by internal reference scaling and Trimmed mean of M normalisation.

### Data analysis

Analyses were performed using R3.6.1. Linear models were generated to estimate the relationship between gene expression and age using the base R function *lm*, which generated p-values indicating the significance of the relationship. Models with a p-value <0.05 were considered significant. Plots were generated using the R package *ggplot2*. Mean protein abundance (for proteomics data) and mean gene expression TPM (for GTEx data) was plotted using the function *ggbarplot*, with the standard error of the mean indicated as error bars. P-values indicating the significance of the difference between gene expression/protein abundance in males and females in each given age group were the result of a Wilcoxon rank sum test for independent groups. For the proteome data, we performed a two-sided student’s t-test. Boxplots comparing gene expression in males and females were generated using the function *ggboxplot*, for which p-values were the result of a Wilcoxon rank sum test for independent groups.

