## Supplements for "COVID-19-related coagulopathy – Is transferrin a missing link?": Figure S1.pdf

ANO6

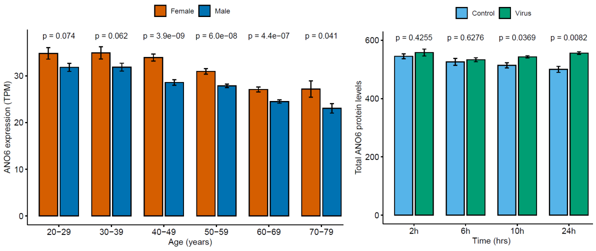


ANXA2

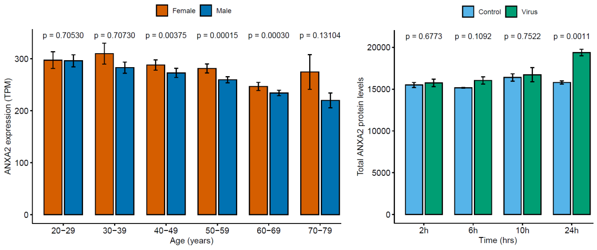


C1QBP

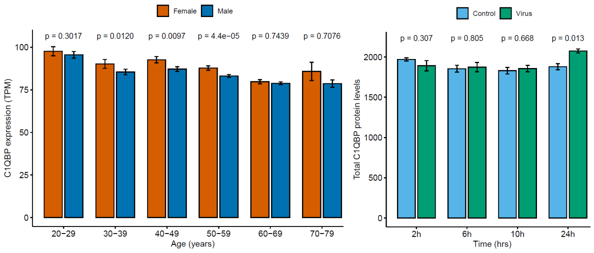


CEACAM1

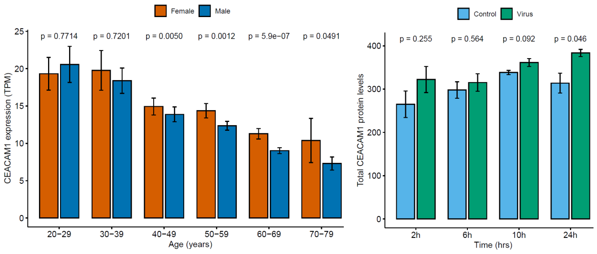


F10

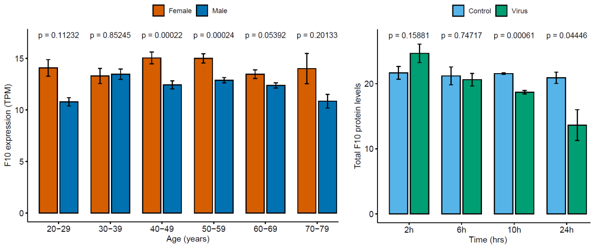


F11R

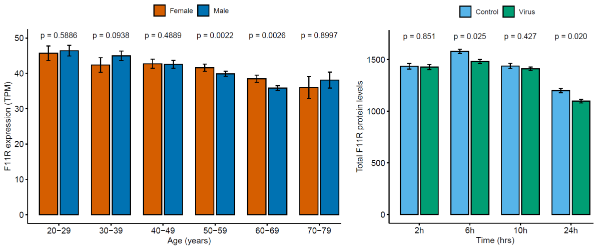


GGCX

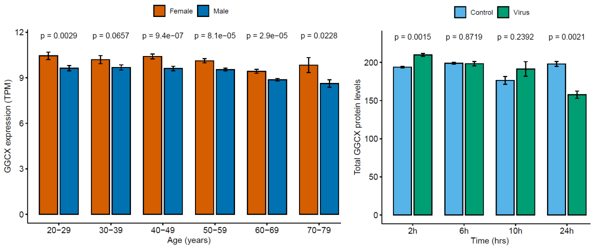


MYH9

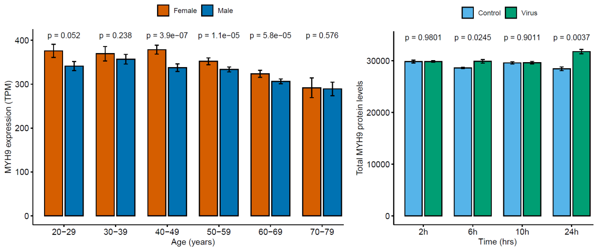


PLA2G4A

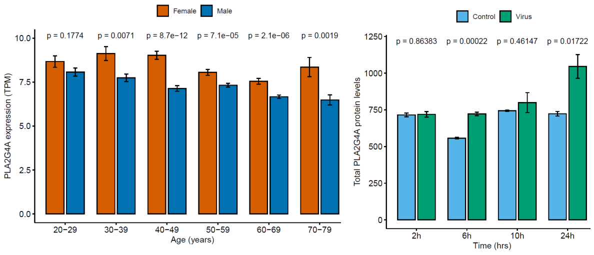

**Figure S1.** Expression of the nine genes associated with the GO term “Blood Coagulation” (GO:0007596) that are differentially regulated (mRNA levels) between females and males, whose expression (mRNA levels) correlates with age (GTEx dataset), and whose expression (protein level) is significantly regulated after SARS-CoV-2 infection. ANXA2 is a coagulation inhibitor, all other proteins are coagulation promoters. Mean gene expression (TPM) is presented across six age groups separately for females and males. P-values were determined using the Wilcoxon rank sum test for independent groups. Moreover, mean protein abundance is shown in uninfected (control) and SARS-CoV-2-infected (virus) Caco-2 cells at different time points post infection. P-values are the result of a two-sided Student’s t-test.
